# Exploring RNA Destabilization Mechanisms in Biomolecular Condensates through Atomistic Simulations

**DOI:** 10.1101/2024.09.13.612876

**Authors:** Matteo Boccalini, Yelyzaveta Berezovska, Giovanni Bussi, Matteo Paloni, Alessandro Barducci

**Affiliations:** Centre de Biologie Structurale (CBS), Université de Montpellier, CNRS, Inserm, Montpellier, France; Molecular and Statistical Biophysics, Scuola Internazionale Superiore di Studi Avanzati, Trieste, Italy; Thomas Young Centre and Department of Chemical Engineering, University College London, London WC1E 7JE, U.K

**Keywords:** RNA Folding, Membraneless Organelles, Molecular Dynamics, Enhanced Sampling, RNA Conformational Ensemble, Protein-RNA Interactions

## Abstract

Biomolecular condensates are currently recognized to play a key role in organizing cellular space and in orchestrating biochemical processes. Despite an increasing interest in characterizing their internal organization at the molecular scale, not much is known about how the densely crowded environment within these condensates affects the structural properties of recruited macromolecules. Here we adopted explicit-solvent all-atom simulations based on a combination of enhanced sampling approaches to investigate how the conformational ensemble of an RNA hairpin is reshaped in a highly-concentrated peptide solution that mimics the interior of a biomolecular condensates. Our simulations indicate that RNA structure is greatly perturbed by this distinctive physico-chemical environment, which weakens RNA secondary structure and promotes extended non-native conformations. The resulting high-resolution picture reveals that RNA unfolding is driven by the effective solvation of nucleobases through hydrogen bonding and stacking interactions with surrounding peptides. This solvent effect can be modulated by the aminoacid composition of the model condensate as proven by the differential RNA behaviour observed in the case of arginine-rich and lysine-rich peptides.

## Introduction

In the cell, RNAs can populate multiple, distinct conformations and this structural plasticity is intimately connected to their biological activities and tightly regulated by specific processes Ken et al. (2023); Cruz and Westhof (2009); Mustoe et al. (2014).

In this respect, it is particularly intriguing to investigate the potential role of biomolecular condensates, which are currently thought to be crucial players in organizing cellular environment Banani et al. (2017). These dynamic non-stoichiometric assemblies are characterized by a high local concentration of specific proteins and nucleic acids and they can act as Membrane-Less Organelles (MLOs) by creating chemically distinct compartments Hyman et al. (2014); Banani et al. (2017); Shin and Brangwynne (2017). The unique physicochemical environment inside MLOs may significantly reshape the conformational landscape of recruited RNA molecules, potentially providing an effective regulatory mechanism Ganser et al. (2019).

Recently, a few pioneering studies have explored this hypothesis by probing the RNA properties in model condensates through controlled *in vitro* experiments Nott et al. (2016); Langdon et al. (2018); Cakmak et al. (2020); Choi et al. (2022); Meyer et al. (2023). While experimental evidence suggests that condensate environment can affect the catalytic activity of ribozymes Poudyal et al. (2019); Iglesias-Artola et al. (2022), RNA structural characterization in biomolecular condensed phase is limited due to the technical difficulties of most structural biology approaches in non-homogeneous and/or highly-concentrated systems Peran and Mittag (2020); Meyer et al. (2023). Förster Resonance Energy Transfer (FRET) spectroscopy has provided some insight on the structure of nucleic acids in model MLOs, showing that *in vitro* condensates can melt duplexes Nott et al. (2016); Choi et al. (2022) and these findings were recently supported by Nuclear Magnetic Resonance (NMR) experiments Rangadurai et al. (2024). Regardless of these inspiring results, a clear picture of RNA structural properties in biomolecular condensates, as well as a detailed description of the molecular interactions that likely reshape the conformational ensemble in this environment, are still lacking.

All-atom Molecular Dynamics (MD) simulations can provide an accurate, high-resolution description of RNA structure and interactions, especially when supplemented with advanced simulation algorithms Bottaro et al. (2016a); Sponer et al. (2018); Mly`nsky` and Bussi (2018); Zerze et al. (2021). Molecular simulations have also played an important role in the investigation of the dynamical and structural properties of biomolecular condensates. While various coarse-grained models Dignon et al. (2018); Choi et al. (2019); Benayad et al. (2020); Nguyen et al. (2022); Joseph et al. (2021); Tesei and Lindorff-Larsen (2022) are often adopted to effectively tackle the length and time scales associated to these large-sized assemblies, a few studies revealed the potential of all-atom MD simulations in this domain Zheng et al. (2020); Galvanetto et al. (2023); Polyansky et al. (2023); Mukherjee and Schäfer (2023). In this respect, we recently developed an efficient, fragment-based strategy to shed light on molecular interactions in condensates by performing all-atom MD simulations of high-concentration mixtures of peptides and/or oligonucleotides Paloni et al. (2020, 2021).

In this work, we built on these results and we extended this computational approach to investigate the conformational landscape of a small structured RNA in a molecular environment closely mimicking the interior of a biomolecular condensate.

## Results

### A minimal system to investigate RNA structures in condensates at the atomistic scale

RNA structure is made up of a limited number of secondary and tertiary motifs, which constitute modular units that form complex architectures Tinoco Jr and Bustamante (1999). Among these units, RNA tetraloops, which display a hairpin structure with Watson-Crick base pairing in the stem region and specific non-canonical pairings in a four-base loop, are ubiquitous in large RNAs Woese et al. (1990) and hence popular experimental and computational models for investigating RNA structure, dynamics, and folding Nozinovic et al. (2010); Tan et al. (2018). Here we chose the GCAA member of the GNRA tetraloop family, whose structure has been solved by NMR and it is characterized by a significant flexibility in the loop region Jucker et al. (1996); Zhao and Xia (2007).

Inspired by previous results Paloni et al. (2020, 2021), we modeled the physico-chemical environment within a biomolecular condensate by simulating the hairpin in a peptide solution with a concentration (around 300 mg/ml) comparable to those observed in model *in vitro* condensates Brady et al. (2017); Murthy et al. (2019); Galvanetto et al. (2023) (Fig. 1b). In particular, we selected a polypeptide sequence (RGRGG) based on the RGG/RG motif, which is frequently involved in RNA binding Thandapani et al. (2013) and it is abundant in low-complexity regions of proteins found in MLOs Chong et al. (2018). Experimental data suggest that individual RGG/RG motifs bind to RNA weakly but they can establish a dynamical network of multivalent interactions when present in multiple copies and ultimately stabilize protein-RNA condensates Takahama et al. (2011); Chong et al. (2018).

**Figure 1.**
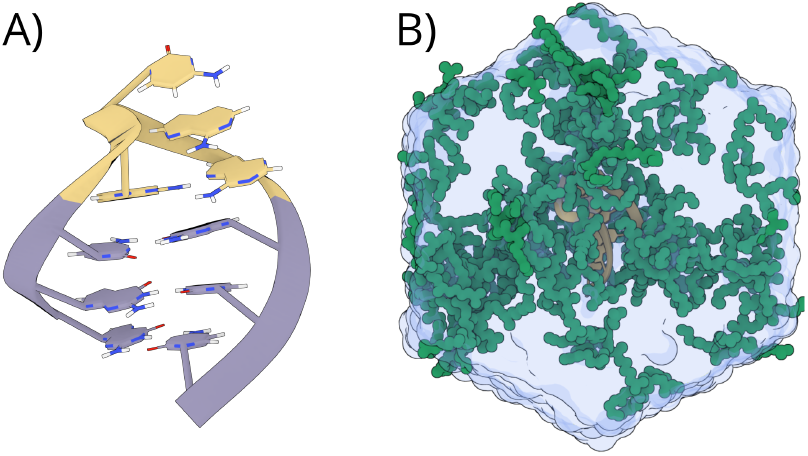
GCAA tetraloop and biomolecular condensate reconstruction: (**A**) Native structure of GCAA tetraloop (pdb-code: 1ZIH); the stem region is represented in purple, the loop region is in yellow. (**B**) Representative snapshot from the MD simulation of GCAA tetraloop (in yellow) within around 300 mg/ml RGRGG (in green) peptide solution that recreate the chemical environment inside biomolecular condensate.

In order to investigate how these intermolecular interactions affect the tetraloop structural properties, we exhaustively sampled its conformational landscape both in the model biomolecular condensed phase and in standard water solution, by combining Well-tempered Metadynamics Barducci et al. (2008); Laio and Parrinello (2002) and Replica Exchange with Solute Tempering Wang et al. (2011). The resulting hybrid protocol (WTMetaD-REST2) has been proven to be particularly effective for characterizing the structural ensemble of small RNA molecules Mlynsky et al. (2022).

### Extended RNA Conformations are favored in biomolecular condensed phase

We report in Fig. 2 the Free-Energy Surfaces (FESs) as a function of structural deviation of the hairpin from the native structure (eRMSD Bottaro et al. (2014)) and its Solvent-Accessible Surface Area (SASA) computed using a standard probe radius. In water (Fig. 2a) the system exhibits a global free-energy minimum at low values of eRMSD and SASA that corresponds to the native ensemble and encompasses available NMR structures (Fig. S3) Jucker et al. (1996), whereas a higher-energy basin representing unfolded states is observed at larger values of eRMSD and SASA. This result suggests that the folded hairpin conformation is stable in standard conditions with a predicted folding free-energy ΔG_folding_ of − 14.1±0.2 kJ/mol, in good agreement with experimental ΔG_folding_ of − 12.6±1.0 kJ/mol SantaLucia Jr et al. (1992) (Fig. S5). This scenario is dramatically altered in the concentrated peptide solution (Fig. 2a). In this environment, the native structure is severely destabilized and the folding landscape is dominated by multiple free-energy minima at higher values of eRMSD. Particularly, the most relevant basin (eRMSD *>* 1.5, SASA ≈ 26) corresponds to unfolded states that are more extended than those observed in water solution.

**Figure 2.**
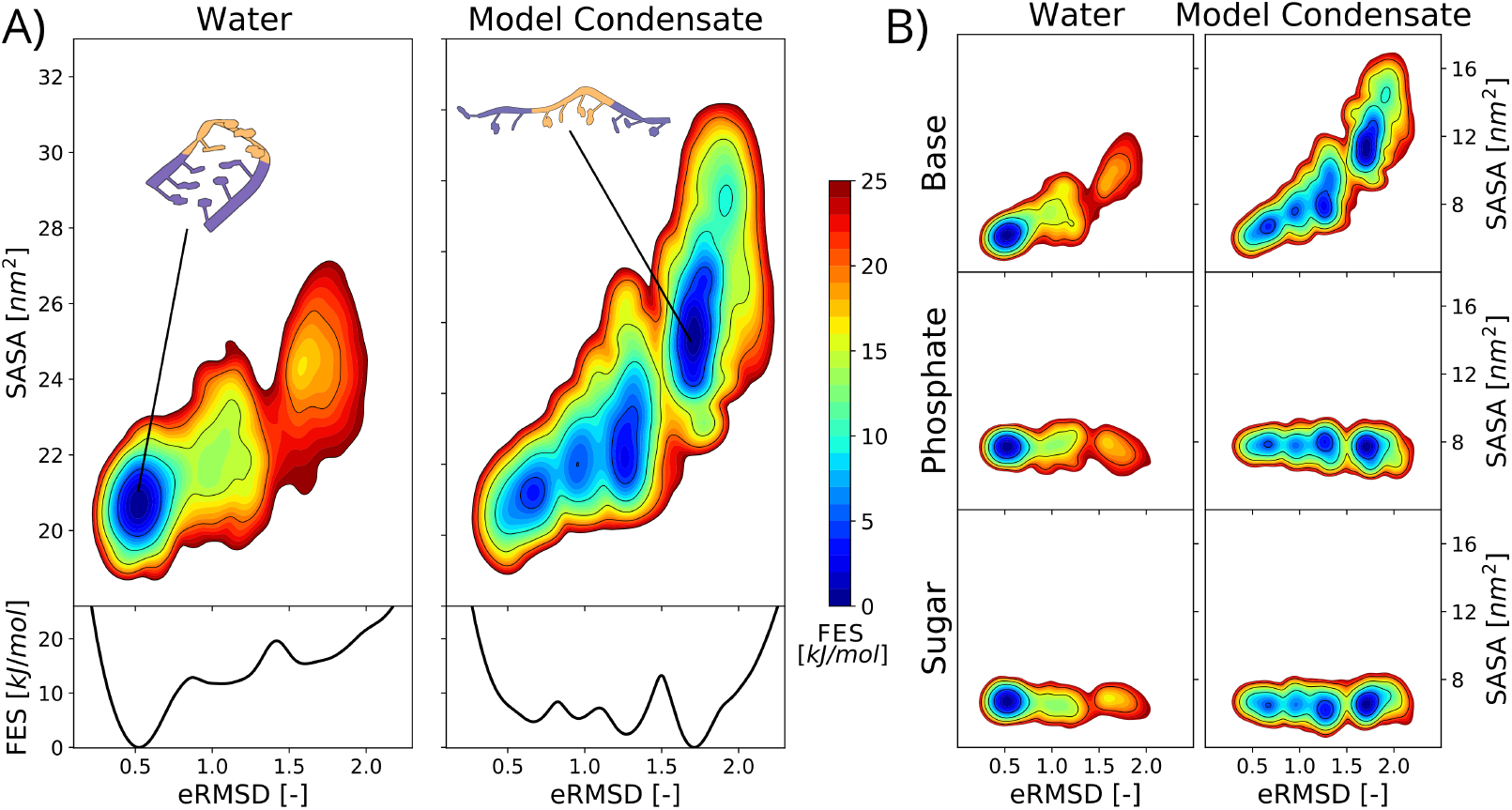
Extended RNA conformations are favored in biomolecular condensed phase. (**A**) Free Energy Surface (FES) as function of eRMSD and Surface Accessible Solvent Area (SASA), obtained from all-atom MD simulation of GCAA tetraloop in water (upper left) and in RGRGG peptide solution (300 mg/ml) mimicking the biomolecular condensate environment (upper right). Molecular representations are reported as an example of GCAA structural conformation corresponding to the main free-energy basin. FES profiles as function of eRMSD for GCAA tetraloop MD simulation in water (lower left) and in model condensate (lower right). (**B**) FES as a function of global eRMSD and SASA related to different RNA functional groups: base (top), phosphate (center), sugar (bottom). The left column is referred to simulation in water while the right one is referred to simulation with model condensate.

This finding suggests that the model condensate represents a better solvent for the RNA molecule leading to its expansion. In order to dissect the role of the different RNA functional groups in this process, we evaluated the free-energy landscape as a function of the eRMSD and the solvent accessibility of phosphate, base or sugar groups (Fig. 2b). The resulting FESs reveal that in both systems the SASA of phosphate and sugar groups is not particularly affected by the unfolding process whereas the nucleobases are remarkably more solvated in unfolded states. This behaviour is particularly pronounced in the biomolecular condensed phase suggesting that the hairpin unfolding in this environment may be attributed to favourable interaction between the RNA bases and the mixed peptide/water solvent.

### Destabilization of RNA Secondary Structures in Condensed Phases

Building on this result, we investigated how intermolecular interactions affect RNA base pairing and destabilize the hairpin secondary structure. We first focused on the three canonical Watson-Crick base pairs that define the stem region of the tetraloop (Fig. 3a). Remarkably, the free-energies of formation of individual pairs (ΔG) indicate that all these native interactions are considerably stable in water whereas they are disfavoured and only transiently observed in the peptide-rich solution. This information can be translated into a more global picture of the folding process by considering the freeenergy as a function of the number of native stem base pairs formed for a given conformation (Nc). In agreement with the analysis reported in 2), the corresponding free-energy profiles (Fig. 3b) confirm that the fullyformed stem (Nc = 3) is the most stable state in water. Conformations with only two base pairings (Nc = 2) are marginally populated while further loss of base-pairing corresponds to large energy penalties and it ultimately leads to a considerable free-energy difference between the folded and the completely open state (Nc = 0). Conversely, base pairing is significantly destabilized in the model condensate. In this environment, the stem can be described with a heterogeneous structural ensemble, where the folded (Nc = 3) and completely unfolded (Nc = 0) states are similarly populated and separated by higher-energy intermediate states (Fig. 3b).

**Figure 3.**
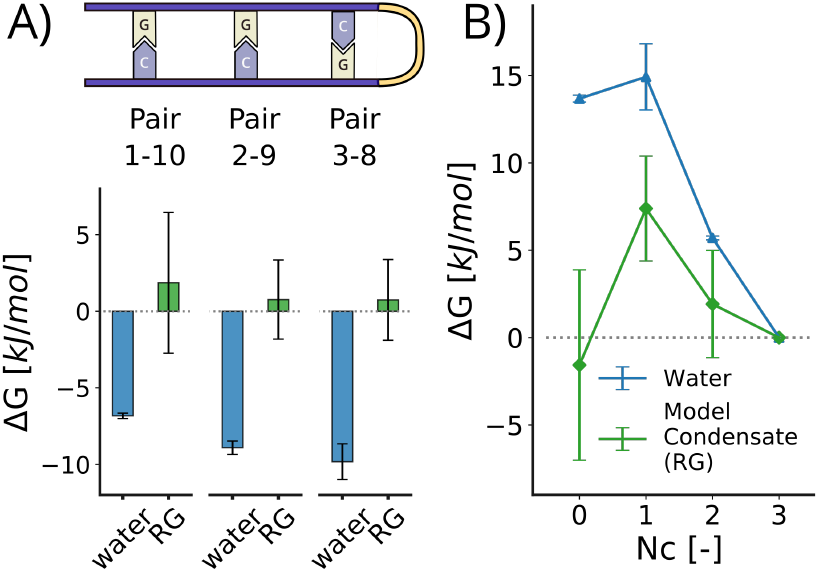
Destabilization of RNA Secondary Structures in Condensed Phases. (**A**) Schematic representation of hairpin stem region (top). Formation free energies (Δ*G*) of individual pairs (G1-C10, G2-C9, C3-G8, respectively). In blue are the values obtained from the simulation of the RNA in water, in green are the values for the RNA within the model condensate consisting of a solution of RGRGG peptides (indicated RG in the figure) (bottom). (**B**) Free-energy (Δ*G*) as a function of the number of native stem base pairs formed (*Nc*). *Nc* = 3 indicates the fully-formed stem while *Nc* = 0 represents the absence of pairing at the stem level. *Nc* = 1 and *Nc* = 2 correspond to an intermediate condition in which the pairing is partially removed. The blue line indicates RNA in water, and the green line refers to RNA within the model condensate (RG).

### Impact of Condensed Phases on RNA Loop Structural Dynamics

Given the major disruption of stable secondary structure in the stem observed in our condensed phase simulations, we then investigated how this environment may affect the structural properties of a flexible region, such as the loop in the GCAA tetraloop Zhao and Xia (2007); Trantírek et al. (2007). To this aim, we limited our analysis to conformations with a fully formed stem (Nc = 3) and we described the structural ensemble of the loop in terms of stacking and pairing interactions among G4-C5-A6-A7 nucleobases.

Our simulations confirm that the loop is structurally flexible in water and displays a variety of transient interactions, including A6-A7 stacking (∼ 90% probability), C5-A6 stacking (∼50%) and the G4-A7 non-canonical pairing (70% of which ∼ 90% trans-Sugar-Hoogsteen pairing) (Fig. 4a left), in very good agreement with NMR-derived conformers Jucker et al. (1996) (Fig. S3) and with analysis of crystallographic structures Bottaro et al. (2016b). This structural ensemble is strongly perturbed in peptide-rich solution where base stacking is significantly weakened and the non-canonical G4-A7 is almost vanished (Fig. 4a right). Remarkably, the model condensate also induced a characteristic, non-native G4-A6 pairing (21% of which ∼ 50% trans-Sugar-Hoogsteen), which is never observed in our simulations in water. We note that this alternative pairing has been suggested as a possible folding intermediate in previous MD simulations performed in water Viegas et al. (2023), although using a different force field.

**Figure 4.**
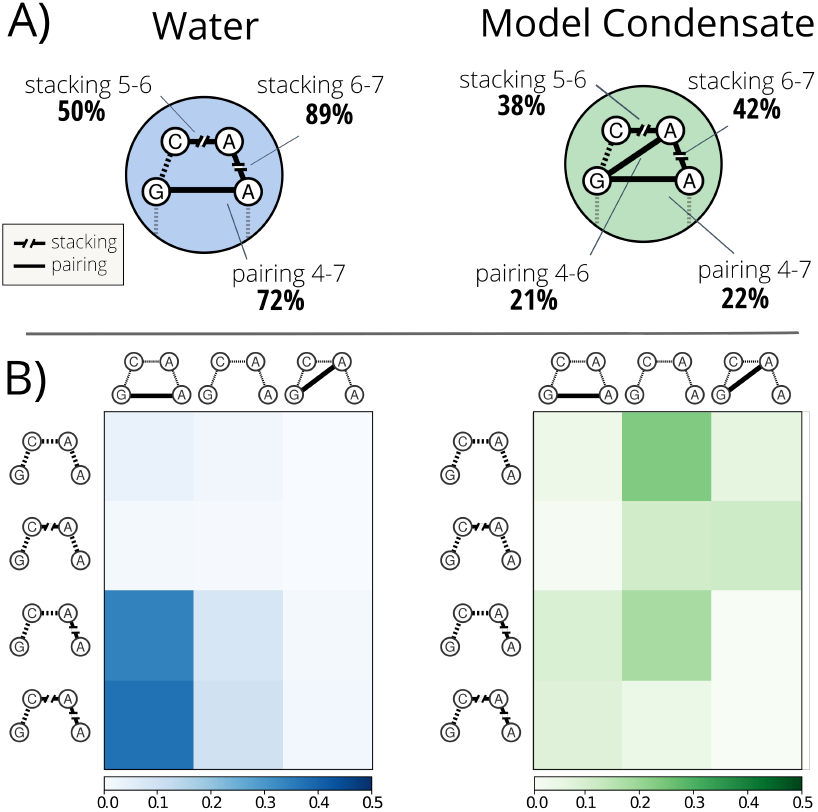
Impact of Condensed Phases on RNA Loop Structural Dynamics. (**A**) Diagram of the most relevant structural features in the loop region. Probabilities were obtained by restricting the analysis to hairpin conformations in which the stem is fully formed (*Nc* = 3). On the left, values for RNA in water, on the right, values for RNA inside the model condensate. (**B**) Probability maps of main loop conformational states in terms of pairing and stacking for structures with fully formed stem (*Nc* = 3). In row, states are distinguished in terms of stacking (from top to bottom: no stacking, C5-A6 stacking, A6-A7 stacking, and C5-A6 stacking + A6-A7 stacking); in column, states are distinguished in terms of pairing (from left to right: G4-A7 pairing, no pairing, G4-A6 pairing). In the left panel (blue) the probabilities of RNA in water, in the right (green) the probabilities of RNA inside the condensed model.

A more global description of the differential behaviour of the loop in the two solvents is provided by the probability maps reported in (Fig. 4b), where we summarized the loop conformational ensemble in 12 discrete states associated to specific stacking and pairing patterns. This analysis confirms that, while our simulations correctly capture the loop flexibility suggested by NMR experiments in water Jucker et al. (1996); Zhao and Xia (2007), they reveal how the condensate environment completely reshapes its conformational ensemble by disfavoring native interactions and possibly promoting non-native pairing.

### Influence of Condensate Composition on RNA Structural Perturbation

Until now we described how our model condensates affect the conformational landscape of GCAA tetraloop, but we have not yet characterized the peptide-RNA interactions that are responsible for the hairpin destabilization. Taking into account the net positive charge of the RGRGG peptide, one can speculate that the electrostatic attraction for the negatively-charged RNA hairpin is sufficient to rationalize the observed effects on its structural properties. In order to test this hypothesis and to shed light on the role of the peptide sequence, we then simulated the GCAA tetraloop in a condensed phase of KGKGG peptides, that display the same positive charge but likely differ in terms of molecular interactions with RNA Paloni et al. (2021). The resulting FES as a function of eRMSD and SASA is reported in Fig. 5 and it shows that while in this condition the GCAA conformational ensemble is significantly reshaped with respect to what observed in water (Fig. 2a), the extent of the condensate-induced perturbation is remarkably smaller than in the case of RGRGG peptides (Fig. 2b). The comparison of hairpin stability in the diverse environments (Fig. 5a bottom panel and Fig. S7), further confirms that the lysine-rich model condensate is only mildly destabilizing and it represents an intermediate scenario between water and arginine-rich model condensate. Analysis of SASA (Fig. 5b) indicates that while RNA expansion is similar in both model condensates, RGRGG peptides have a larger RNA interaction surface acting as a better RNA solvent. This difference has to be traced back to a stronger propensity of RGRGG peptides to interact with RNA and to induce its unfolding that cannot be rationalized in terms of net electrostatic attraction. We therefore analyzed the H-bonding between peptide side chains and the diverse RNA chemical groups by separately considering the conformations in which the RNA is folded (eRMSD *<* 1.5) and unfolded (eRMSD *>* 1.5) (Fig. 5c). We first note that overall RGRGG peptides form more H-bonds than KGKGG ones with all chemical groups. Nevertheless, the number of peptide-RNA H-bonds seems not to be affected by RNA (un)folding in most of the cases. Strikingly, the only noticeable exception is observed for arginine-nucleobases H-bonds that significantly increase upon hairpin unfolding. This behaviour is not observed in lysine-rich model condensate suggesting a distinctive capability of arginine side chains to strongly interact with unpaired nucleobases, eventually inducing a stronger perturbation on RNA free energy landscape. Furthermore, we expect that this differential destabilization is further reinforced by energetically-favourable stacking interactions between nucleobases and arginine guanidinium groups, which preferentially takes place in RNA unfolded states (Fig. 5d).

**Figure 5.**
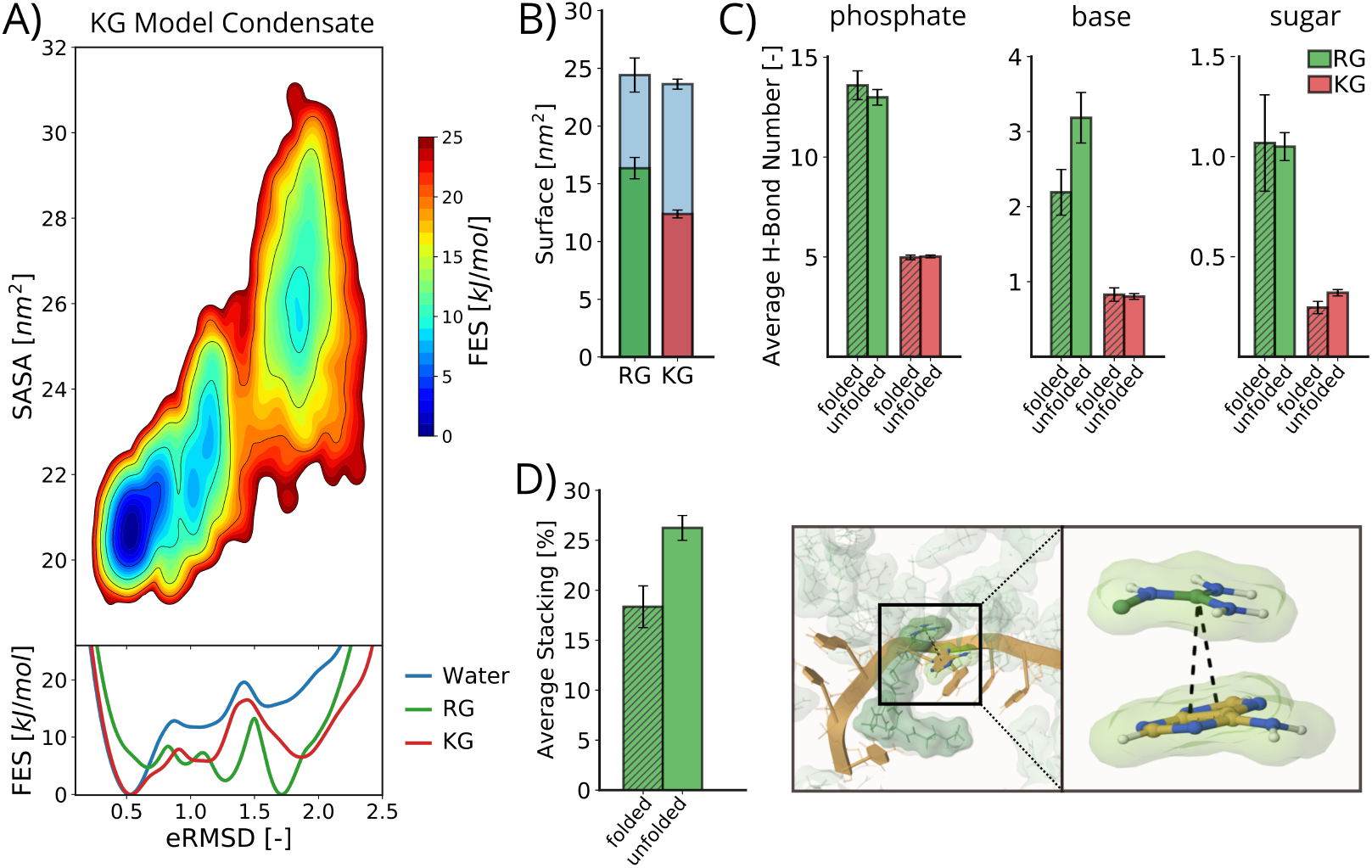
Influence of condensate composition on RNA structural perturbation. (**A**) Free Energy Surface (FES) as function of eRMSD and Surface Accessible Solvent Area (SASA), obtained from simulation of GCAA tetraloop in KGKGG peptide solution (KG model condensate) (top). FES profiles as function of eRMSD for GCAA tetraloop simulation in water (blue), RG model condensate (green) and KG model condensate (red) (bottom). (**B**) In blue are reported tetraloop SASA in RG model condensate (left) and in KG model condensate (right). In green is represented the RNA-peptides interaction surface in RG model condensate (left) while in red the RNA-peptides interaction surface in KG model condensate (right). (**C**) Average hydrogen bond number between RNA and peptide side chains, reported for different RNA functional groups: phosphate (left), base (center), sugar (right). Hatched bars are referred to folded hairpin conformations (*eRMSD <* 1.5) while unhatched bars are referred to unfolded conformations (*eRMSD >* 1.5). In green RG model condensate while in red KG model condensate. (**D**) Average percentage of hairpin nuclobases forming intermolecular stacking interactions with the arginine guanidinium group in RG model condensate. The hatched bar is referred to folded hairpin conformations (*eRMSD <* 1.5) while unhatched bar is referred to unfolded conformations (*eRMSD >* 1.5). On the right a molecular representation of intermolecular stacking interaction between GCAA nucleobase and guanidinium group.

## Discussion

Biomolecular condensates have recently emerged as a fundamental organizing principle of the cellular space Hyman et al. (2014); Banani et al. (2017); Shin and Brangwynne (2017). Even if this field of research is still in its infancy and contrasting hypotheses have been proposed about the molecular mechanism of condensate assembly, an increasing body of evidence suggests that biomolecular condensation is a key process for regulating RNA activity inside the cell Wiedner and Giudice (2021); Ripin and Parker (2023).

Quantitative characterization of this regulation process inherently requires understanding how the peculiar environment of biomolecular condensed phases can affect the conformational ensemble of recruited RNA molecules. In order to shed some light on this topic, we used explicit-solvent all-atom MD simulations to investigate the folding landscape of an RNA tetraloop in a highly-concentrated solution of an arginine-rich peptide, which mimics the interior of a biomolecular condensate. To this aim, we relied on an efficient enhanced sampling method based on the combination of WT-MetaD Barducci et al. (2008); Laio and Parrinello (2002) and Replica-Exchange Solute Tempering method (REST2) Wang et al. (2011) that has been successfully applied to determine the conformational landscape of several RNA hairpins in standard conditions Mlynsky et al. (2022). Similarly, we first took advantage of this computational strategy to predict a folding free-energy landscape of the GCAA tetraloop in water solution. Our results are in good agreement with previous MD simulations Tan et al. (2018); Zerze et al. (2021) performed using a very similar force field but different enhanced sampling methods, confirming the effectiveness of REST2-MetaD protocol. Importantly, our results are also in good agreement with available NMR structures Jucker et al. (1996) and thermodynamic data from calorimetry experiments SantaLucia Jr et al. (1992); Dale et al. (2000), thus confirming the accuracy of last-generation RNA force fields Tucker et al. (2022) for this class of systems.

Most importantly, we then employed this advanced simulation scheme to determine the free-energy landscape of the system in our model biomolecular condensed phase, thus providing an unprecedented picture of RNA conformational ensemble in these extreme conditions. Our simulations revealed that the condensate environment completely reshapes the FES of the GCAA tetraloop leading to a rugged surface (Fig. 1) with multiple basins, where the native structure does no longer represent the global free-energy minimum. The resulting heterogeneous landscape is in good agreement with FRET experiments suggesting a significant structural disorder and a rich conformational dynamics for small single-stranded RNA molecules in LAF-1 condensates Elbaum-Garfinkle et al. (2015); Kim and Myong (2016). Furthermore, our simulations indicate that direct interactions between the peptides and the RNA bases greatly destabilize the RNA secondary structure and ultimately favor open, unfolded conformations. This destabilization of RNA base pairing is compatible with multiple experimental findings, such as the melting of nucleic acid duplexes in model condensate of disordered N-terminus of DDX4 proteins Nott et al. (2016) or the disruption of the s2m RNA stem-loop structure within a condensate of the SARS-CoV-2 nucleocapsid protein de Vries et al. (2024). Beyond the overall destabilization, our simulations reveal the partial stabilization of alternative, non-native base pairing in the flexible loop of the GCAA hairpin. While the extent of this phenomenon is limited in the model system investigated here, this finding supports the fascinating hypothesis that cellular condensates may induce alternative functionally-relevant structural arrangements in larger RNA molecules. Even if present computational resources do not allow to characterize the structural ensemble of complex cellular RNAs by atomistic MD, the development of CG models for RNA in condensates Nguyen et al. (2022); Yasuda et al. (2025) may provide a complementary approach to tackle this challenge.

Overall, the structural perturbation and conformational redistribution effects are likely tuned by the chemical composition of the condensate. In order to shed more light on this topic, we tested the hairpin behavior in a similar yet distinct environment by substituting arginine with lysine residues in the peptide sequence. The resulting FES indicates that lysine-rich peptides are less destabilizing and lead to a smaller perturbation of the RNA conformational ensemble, in agreement with the experimental observation that polyR/polyD coacervates dissolve RNA duplexes whereas this “helicase activity” is less evident for polyK/polyD condensates Choi et al. (2022). In this respect, our results indicate that arginine side chains interact more effectively with RNA than lysine. In particular, the guanidinium groups can establish a higher number of hydrogen bonds with phosphate, sugar, and nucleobases. In the context of RNA conformational dynamics, arginine-base interactions are particularly relevant since they significantly increase upon RNA unfolding and remarkably stabilizing extended conformations, at variance with what is observed for lysine. This differential destabilizing behaviour is further reinforced by arginine propensity of forming stacking interactions with RNA bases with a mechanism reminiscent of what has been proposed to rationalize urea-driven denaturation of structured RNAs Priyakumar et al. (2009).

Overall, our work shows that biomolecular condensates may play a central role in regulating the biological activity of RNA within the cell by altering their conformational ensemble. In condensed phase, the dynamic network of intermolecular interactions can significantly perturb conformational distribution and affect energy barriers possibly favoring conformational excited states with functional relevance. Moreover, dysregulation of RNA structural ensembles has been linked to various human diseases Bernat and Disney (2015); Conti and Oppikofer (2022); Bose et al. (2024). In this context, elucidating the regulatory role of biomolecular condensates in RNA conformational dynamics could provide insight into the molecular basis of these conditions and reveal potential therapeutic targets for RNA-associated pathologies.

## Methods

MD simulations were performed using GROMACS 2021.4 Abraham et al. (2015) patched with PLUMED 2.8 Tribello et al. (2014); consortium (2019) to perform Well-Tempered Metadynamics Barducci et al. (2008) (WT-MetaD) and using a general Hamiltonian replica exchange implementation Bussi (2014). The initial conformation of the (GGCGCAAGCC) RNA tetraloop was the model 1 of NMR structures deposited as Protein Data Bank (PDB) entry from 1ZIH pdb Jucker et al. (1996). The GCAA tetraloop was modeled with the DES-Amber force field developed by Shaw and co-workers Tucker et al. (2022). In the REST2 Wang et al. (2011) frame-work, 16 replicas were simulated in the effective temperature range 300 - 700 K (using scaling factors in a geometric series between from 1.0 to 0.414). As collective biases for WT-MetaD Barducci et al. (2008) were used in combination eRMSD metric Bottaro et al. (2014) and the gyration radius calculated over all the heavy atoms of RNA backbone 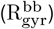. Further details about the simulation protocol and analysis methods are available in the SI.

## Data Availability

Data to reproduce the results of this study (including simulation trajectories) have been deposited on Zenodo (10.5281/zenodo.14203857).

## Acknowledgments

We acknowledge the support of the Swiss National Science Foundation (SNSF) under grant CRSII5_-193740 and of the French Agence Nationale de la Recherche (ANR) under grant ANR-21-CE30-0001. M.B. was supported by funds of the LabMuse EpiGen-Med. G.B. acknowledges the Italian National Centre for HPC, Big Data, and Quantum Computing (grant No. CN00000013), funded within the Next Generation EU initiative. This work was granted access to the HPC resources of CINES under the allocation 2021-A0100712465 made by GENCI.

