## Supplemental Information for "Exploring RNA Destabilization Mechanisms in Biomolecular Condensates through Atomistic Simulations"

### Supplementary Information

#### Simulation protocol

MD simulation were performed using GROMACS 2021.4 Abraham et al. (2015) patched with PLUMED 2.8 Tribello et al. (2014); Plumed-Consortium (2019) to perform Well-Tempered Metadynamics Barducci et al. (2008) (WT-MetaD) and using a general Hamiltonian replica exchange implementation Bussi (2014). The initial conformation of the (GGCGCAAGCC) RNA tetraloop was the model 1 of NMR structures deposited as Protein Data Bank (PDB) entry from 1ZIH pdb Jucker et al. (1996). The GCAA tetraloop was modeled with the DES-Amber force field developed by Shaw and co-workers Tucker et al. (2022). The starting configuration was initially solvated in a rhombic dodecahedron box volume of $\sim 220\ nm^3$. In case of model condensate simulation, peptides (RGRGG or KGKGG) were added inside the box ($\sim 300$ mg/ml) using GMX INSERT-MOLECULES tool. RGRGG and KGKGG structures were created with LEaP program in AmberTools18 Case et al. (2005) and capped with ACE and NME groups to avoid artificial effects induced by the charged termini. The simulation box was neutralized with NaCl, adjusting the salt concentration to 0.1 M. A first 100 ps equilibration was run in NVT ensemble (T=300 K) followed by 100 ps NPT simulation (T=300 K, P = 1 bar) controlling temperature and pressure respectively by means of the v-rescale Bussi et al. (2007) and the Parrinello-Rahman Parrinello and Rahman (1981) schemes. For all simulations, periodic boundary conditions were applied, and long-range electrostatic interactions were evaluated by using the particle mesh Ewald algorithm Essmann et al. (1995) with a cutoff of 1 nm for the real space interactions, while van der Waals interactions were computed by using a cutoff distance of 1 nm. All bond lengths were constrained by using LINCS Hess et al. (1997), allowing for a time step of 2 fs for the integration of the equations of motion. The GCAA tetraloop production-biased simulation in water,in RGRGG and in KGKGG solution were run for 3 $\mu s$, 6 $\mu s$, 5 $\mu s$ respectively. Before starting the production-biased simulation, a REST2 simulation was running for 100 ns.

#### Enhanced Sampling Protocol

In the REST2 Wang et al. (2011) framework, 16 replicas were simulated in the effective temperature range 300 - 700 K (using scaling factors in a geometric series between from 1.0 to 0.414) and replica exchanges were attempted every 1000 steps. The eRMSD metric Bottaro et al. (2014) and the gyration radius calculated over all the heavy atoms of RNA backbone ($\mathrm{R}_{\mathrm{gyr}}^{\mathrm{bb}}$) were used as collective variables for WT-MetaD Barducci et al. (2008); Laio and Parrinello (2002). For the calculation of the eRMSD, the initial NMR structure was used as reference and the cutoff distance was set to 3.2, as it has been shown to be an appropriate choice in the metaD context Bottaro et al. (2016). The metaD bias was accumulated by depositing a gaussian every 1 ps (500 steps); the Gaussian width ($\sigma$ parameter) was set 0.05 and 0.02 for $\mathrm{eRMSD}_{\mathrm{MetaD}}$ and $\mathrm{R}_{\mathrm{gyr}}^{\mathrm{bb}}$ respectively. The initial Gaussian height was of 0.5 kJ/mol with a bias factor of 15. Harmonic restraints potentials were applied to limit the sampling within the interval 0.6 and 1.7 of $\mathrm{R}_{\mathrm{gyr}}^{\mathrm{bb}}$ with a force constant of 1250 kJ/mol, using the LOWER_WALLS and UPPER_WALLS biases available in PLUMED 2.8.

#### Analysis Methods

RNA eRMSD Bottaro et al. (2014) in the analysis ($\mathrm{eRMSD}_{\mathrm{analysis}}$) was computed using PLUMED 2.8 Tribello et al. (2014); the first NMR structure (1ZIH) was used as reference with a cutoff distance of 2.4. All GROMACS analysis (GMX commands) reported below were performed with GROMACS 2021.4 Abraham et al. (2015). RNA Solvent Accessible Surface Areas (SASA) were computed using GMX SASA tool with a solvent probe of 0.14 nm radius; surfaces of interaction between RNA and peptide solvent were obtained as the difference between the sum of the exposed RNA and peptides surfaces (considered independently) and the total surface area of the two species grouped together. Intramolecular RNA pairing and base stacking was evaluated using BARNABA software Bottaro et al. (2019). Hydrogen bond analysis between RNA groups and peptides (considering separately main-chains and side-chains) was performed with the GMX HBOND tool using the default definition of H-bond. The guanidinium group of arginine and nucleobases were considered to be forming a stacking interaction when the distance between the CZ atom of arginine and the center of geometry of the base ring was lower than 0.6 nm (GMX PAIRDIST), and the angle between the plane of the guanidinium group and that of the base is lower than 30 (GMX GANGLE) Marsili et al. (2008); Paloni et al. (2020). Unbiased probability distributions of analyzed quantities were obtained by reweighting the statistics accumulated in REST2-MetaD trajectories using final time-averaged bias potentials deposited on the $\mathrm{eRMSD}_{\mathrm{MetaD}}$ and $\mathrm{R}_{\mathrm{gyr}}^{\mathrm{bb}}$ coordinates. Free energy surfaces (FESs) were computed with the standard equation $F(x) = -k_B T ln(P(x)) + C$, where P(x) is the unbiased probability density, and C is an immaterial constant. Trajectory snapshots were visualized using *Mol** (molstar) toolkit Sehnal et al. (2021). Schematic pictures and panel figures editing was realized using Inkscape Inkscape Project (2020), open-source vector graphics editor.

##### Error estimation

Error bars were obtained by dividing all production-biased simulations in sub-trajectories of $1 \mu s$ long. All sub-trajectories were reweighted using the corresponding final time-average bias obtained from MetaD gaussian deposition. The error is estimated as the standard deviation of values obtained from the different $1 \mu s$ long trajectories.

#### Supplementary Information - Bibliography


Abraham, M. J., Murtola, T., Schulz, R., Páll, S., Smith, J. C., Hess, B., and Lindahl, E. (2015). Gromacs: High performance molecular simulations through multi-level parallelism from laptops to supercomputers. *SoftwareX*, 1:19–25.

Barducci, A., Bussi, G., and Parrinello, M. (2008). Well-tempered metadynamics: a smoothly converging and tunable free-energy method. *Physical review letters*, 100(2):020603.

Bottaro, S., Di Palma, F., and Bussi, G. (2014). The role of nucleobase interactions in rna structure and dynamics. *Nucleic acids research*, 42(21):13306–13314.

Bottaro, S., Banas, P., Sponer, J., and Bussi, G. (2016). Free energy landscape of gaga and uucg rna tetraloops. *The journal of physical chemistry letters*, 7(20):4032–4038.

Bottaro, S., Bussi, G., Pinamonti, G., Reißer, S., Boomsma, W., and Lindorff-Larsen, K. (2019). Barnaba: software for analysis of nucleic acid structures and trajectories. *RNA*, 25(2):219–231.

Bussi, G. (2014). Hamiltonian replica exchange in gromacs: a flexible implementation. *Molecular Physics*, 112(3-4):379–384.

Bussi, G., Donadio, D., and Parrinello, M. (2007). Canonical sampling through velocity rescaling. *The Journal of chemical physics*, 126(1).

Case, D. A., Cheatham III, T. E., Darden, T., Gohlke, H., Luo, R., Merz Jr, K. M., Onufriev, A., Simmerling, C., Wang, B., and Woods, R. J. (2005). The amber biomolecular simulation programs. *Journal of computational chemistry*, 26(16):1668–1688.

Dale, T., Smith, R., and Serra, M. J. (2000). A test of the model to predict unusually stable rna hairpin loop stability. *Rna*, 6(4):608–615.

Essmann, U., Perera, L., Berkowitz, M. L., Darden, T., Lee, H., and Pedersen, L. G. (1995). A smooth particle mesh ewald method. *The Journal of chemical physics*, 103(19):8577–8593.

Hess, B., Bekker, H., Berendsen, H. J., and Fraaije, J. G. (1997). Lincs: A linear constraint solver for molecular simulations. *Journal of computational chemistry*, 18(12):1463–1472.

Inkscape Project. Inkscape, (2020).

Jucker, F. M., Heus, H. A., Yip, P. F., Moors, E. H., and Pardi, A. (1996). A network of heterogeneous hydrogen bonds in gnra tetraloops. *Journal of molecular biology*, 264(5):968–980.

Laio, A. and Parrinello, M. (2002). Escaping free-energy minima. *Proceedings of the national academy of sciences*, 99(20):12562–12566.

Lorenz, R., Bernhart, S. H., Höner zu Siederdissen, C., Tafer, H., Flamm, C., Stadler, P. F., and Hofacker, I. L. (2011). Viennarna package 2.0. *Algorithms for molecular biology*, 6:1–14.

Marsili, S., Chelli, R., Schettino, V., and Procacci, P. (2008). Thermodynamics of stacking interactions in proteins. *Physical Chemistry Chemical Physics*, 10(19):2673–2685.

Paloni, M., Bailly, R., Ciandrini, L., and Barducci, A. (2020). Unraveling molecular interactions in liquid–liquid phase separation of disordered proteins by atomistic simulations. *The Journal of Physical Chemistry B*, 124(41):9009–9016.

Parrinello, M. and Rahman, A. (1981). Polymorphic transitions in single crystals: A new molecular dynamics method. *Journal of Applied physics*, 52(12):7182–7190.

Plumed-Consortium. (2019). Promoting transparency and reproducibility in enhanced molecular simulations. *Nature methods*, 16(8):670–673.

SantaLucia Jr, J., Kierzek, R., and Turner, D. H. (1992). Context dependence of hydrogen bond free energy revealed by substitutions in an rna hairpin. *Science*, 256(5054):217–219.

Sehnal, D., Bittrich, S., Deshpande, M., Svobodová, R., Berka, K., Bazgier, V., Velankar, S., Burley, S. K., Koča, J., and Rose, A. S. (2021). Mol* viewer: modern web app for 3d visualization and analysis of large biomolecular structures. *Nucleic acids research*, 49(W1):W431–W437.

Tribello, G. A., Bonomi, M., Branduardi, D., Camilloni, C., and Bussi, G. (2014). Plumed 2: New feathers for an old bird. *Computer physics communications*, 185(2):604–613.

Tucker, M. R., Piana, S., Tan, D., LeVine, M. V., and Shaw, D. E. (2022). Development of force field parameters for the simulation of single-and double-stranded dna molecules and dna–protein complexes. *The Journal of Physical Chemistry B*, 126(24):4442–4457.

Wang, L., Friesner, R. A., and Berne, B. (2011). Replica exchange with solute scaling: a more efficient version of replica exchange with solute tempering (rest2). *The Journal of Physical Chemistry B*, 115(30):9431–9438.

#### Supplementary Information - Figures

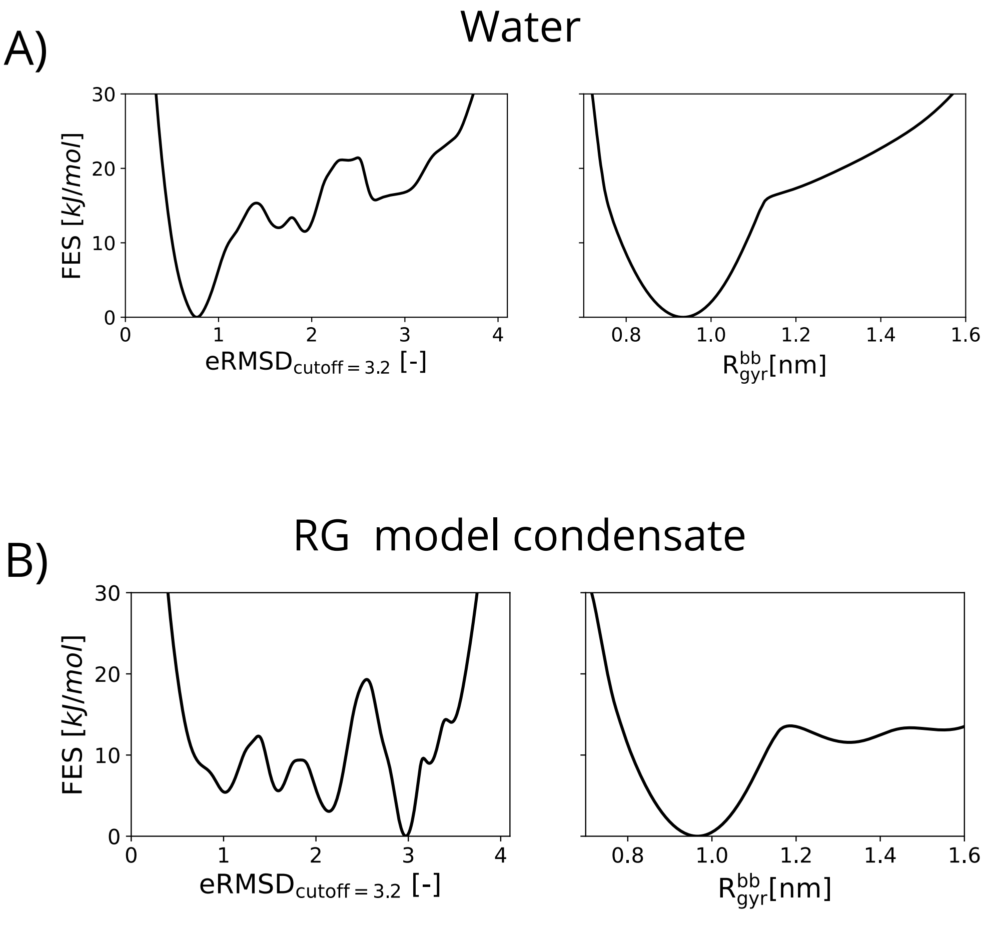


**Figure S1.** Free energy profiles along WT-MetaD Collective Variables (CVs). (**A**) Free energy as a function of eRMSD from the NMR structure (pdb:1ZIH; model=1) with a cutoff=3.2 (left) and the gyration radius calculated over all the heavy atoms of RNA backbone ($R^{bb}_{gyr}$) (right) for GCAA tetraloop simulation in water. (**B**) Free energy as a function of eRMSD from the NMR structure (pdb:1ZIH; model=1) with a cutoff=3.2 (left) and the gyration radius calculated over all the heavy atoms of RNA backbone ($R^{bb}_{gyr}$) (right) for GCAA tetraloop simulation in RG model condensate (RGRGG $\sim$ 300 mg/ml ).

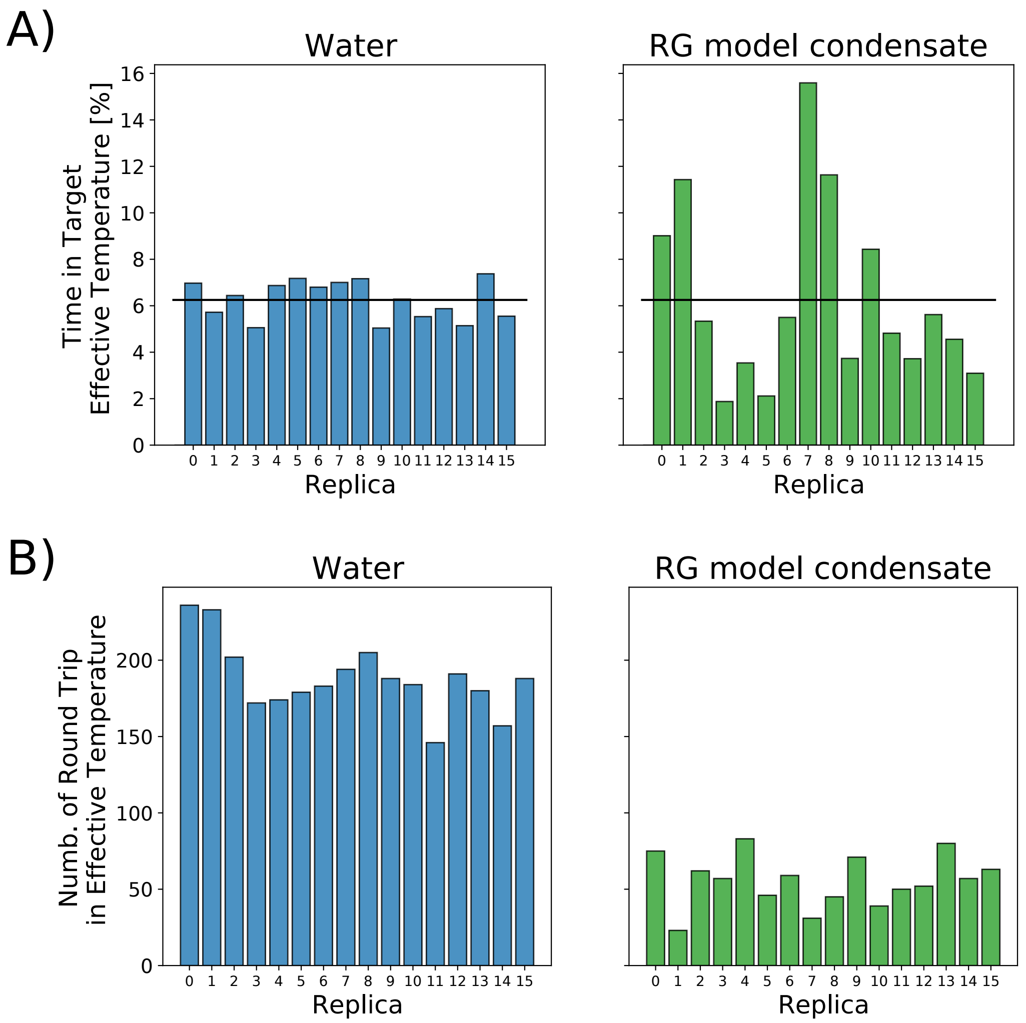


**Figure S2.** Diffusion of replicas in effective temperature space. (**A**) Percentage of time spent in target effective temperature for different replica. In blue (left) are reported data for GCAA tetraloop in water while in green (right) data for GCAA tetraloop in RG model condensate; the black line shows the ideal percentage of 6.25% in which all the replica spend the same time in each effective temperature. (**B**) Number of round trip in effective temperature. The round trip for a replica refers to travel from the lowest temperature to the highest and back. In blue (left) are reported data for GCAA tetraloop in water while in green (right) data for GCAA tetraloop in RG model condensate.

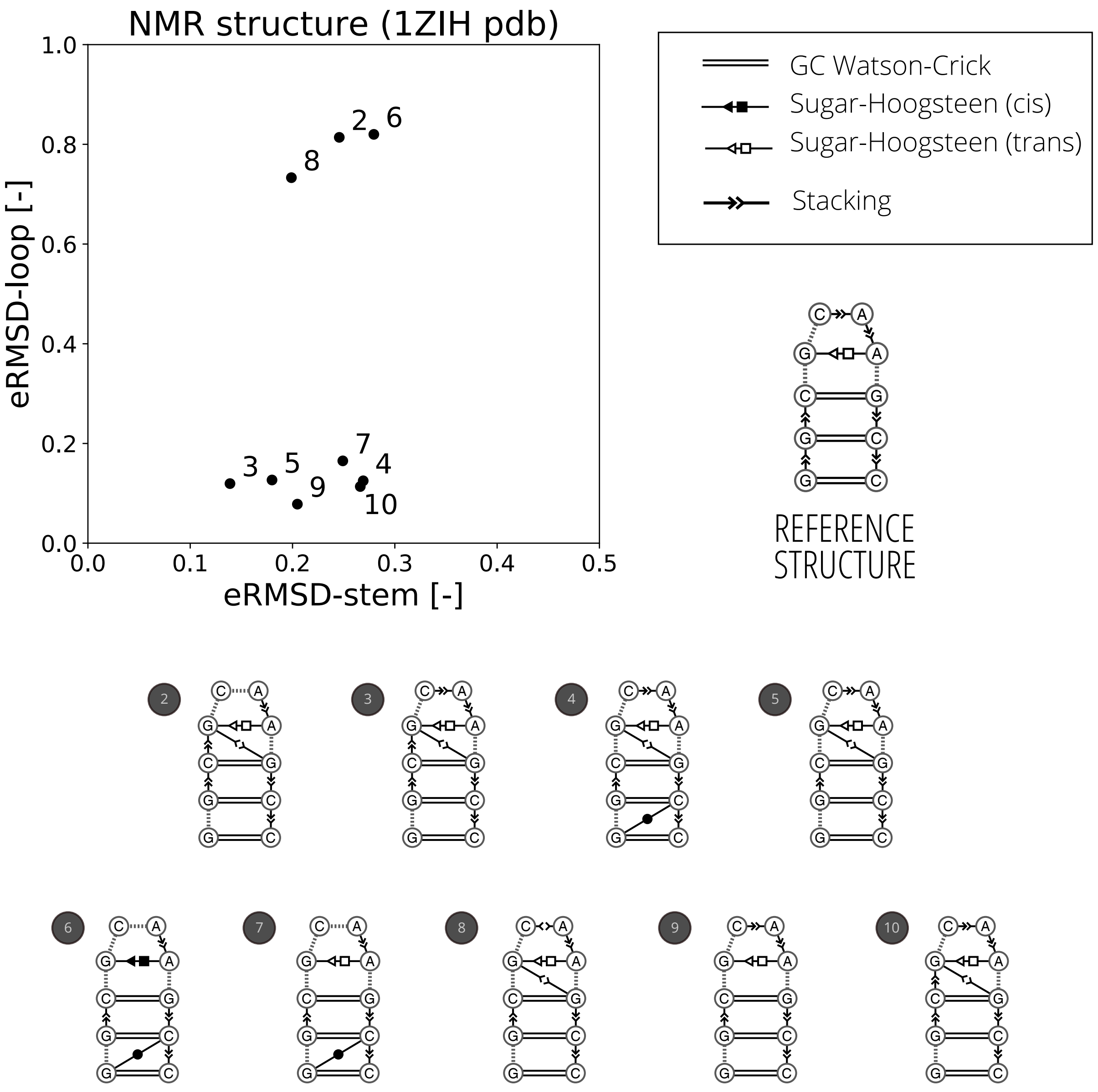


**Figure S3.** GCAA conformers in 1ZIH pdb Jucker et al. (1996). Secondary structure representation (with Leontis–Westhof classification) of GCAA structures in pdb:1ZIH obtained with Barnaba software Bottaro et al. (2019). The reference structure in eRMSD calculation corresponds to model 1 in pdb:1ZIH. In the top part of the panel all GCAA conformers are represented in the $\mathrm{eRMSD}_{\mathrm{stem}}$ and $\mathrm{eRMSD}_{\mathrm{loop}}$ space. $\mathrm{eRMSD}_{\mathrm{stem}}$ is calculated using as reference structure the stem region of the reference conformer (residues G1, G2, C3, G8, C9, C10) while $\mathrm{eRMSD}_{\mathrm{loop}}$ is calculated using the loop of the reference conformer (G4, C5, A6, A7). In the scatter plot, we can observe that the stem is stable ( all conformers $\mathrm{eRMSD}_{\mathrm{stem}} < 0.3$ ) while the loop exhibits a larger heterogeneity ($\mathrm{eRMSD}_{\mathrm{loop}} > 0.6$ for conformers 2,6,8).

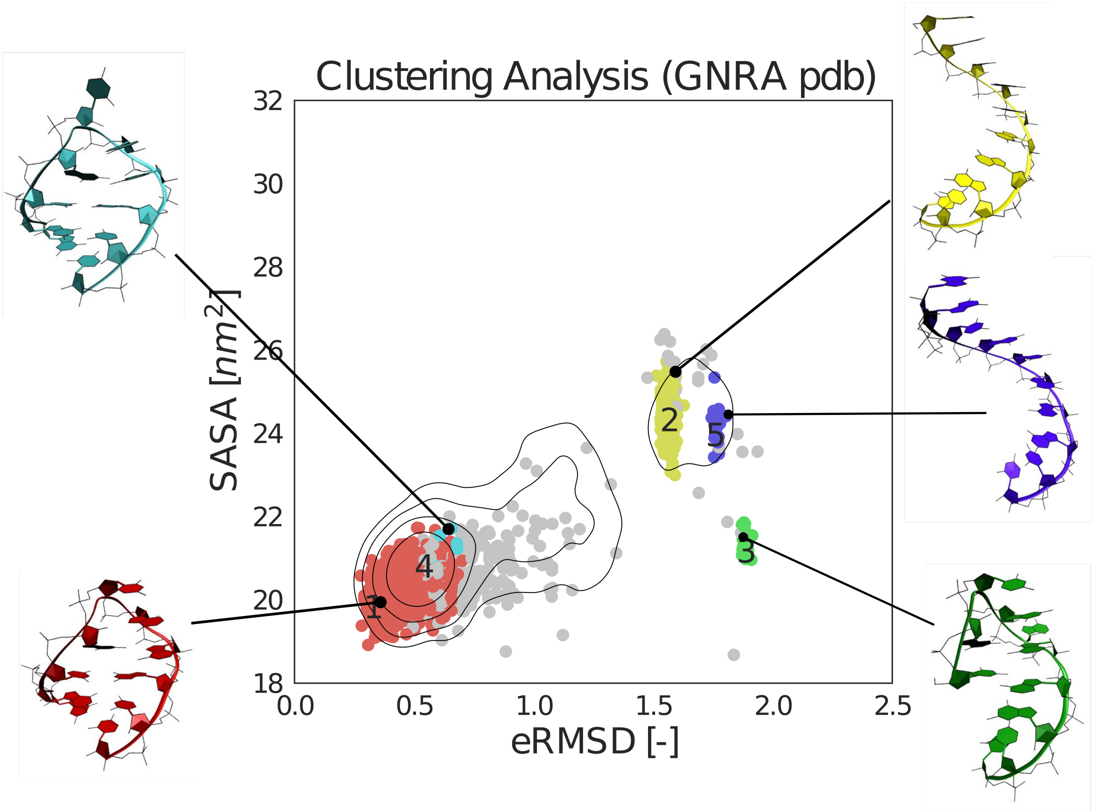


**Figure S4.** Cluster analysis of GNRA tetraloop structure in pdb database. In figure are reported GNRA tetraloop structures (in which the N could be either uracil, adenine, cytosine, or guanine, and the R is either guanine or adenine) projected on the eRMSD and SASA space. The contour plot represents the free energy surface for GCAA tetraloop simulation in water. Principal clusters of GNRA in pdb database are colored in the picture and the corresponding structure representations (centers of the clusters), colored as in the plot, are reported. For cluster analysis it has been used Barnaba software Bottaro et al. (2019).

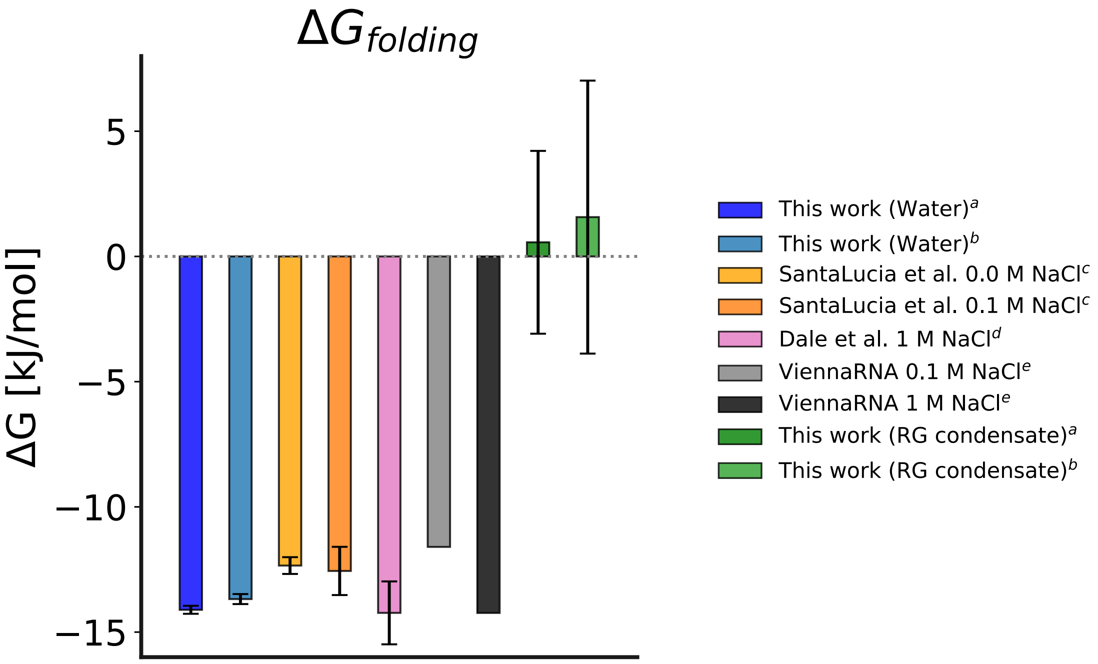


**Figure S5.** Folding free energy of GCAA tetraloop. In blue are reported the folding free energy calculated from GCAA simulation in water and in green for GCAA simulation in RG-model condensate. In both cases, two different methods (a) and (b) have been used.

(**a**) Folding free energy estimated as free energy differences between folded and unfolded conformation individuated using a threshold on eRMSD (eRMSD = 1.5).

(**b**) Folding free energy estimated as free energy differences between folded and unfolded conformation based on secondary structure (folded: $N_c = 3$; unfolded: $N_c = 0$ ).

(**c**) SantaLucia et al. SantaLucia Jr et al. (1992)

(**d**) Dale et al. Dale et al. (2000)

(**e**) ViennaRNA Lorenz et al. (2011)

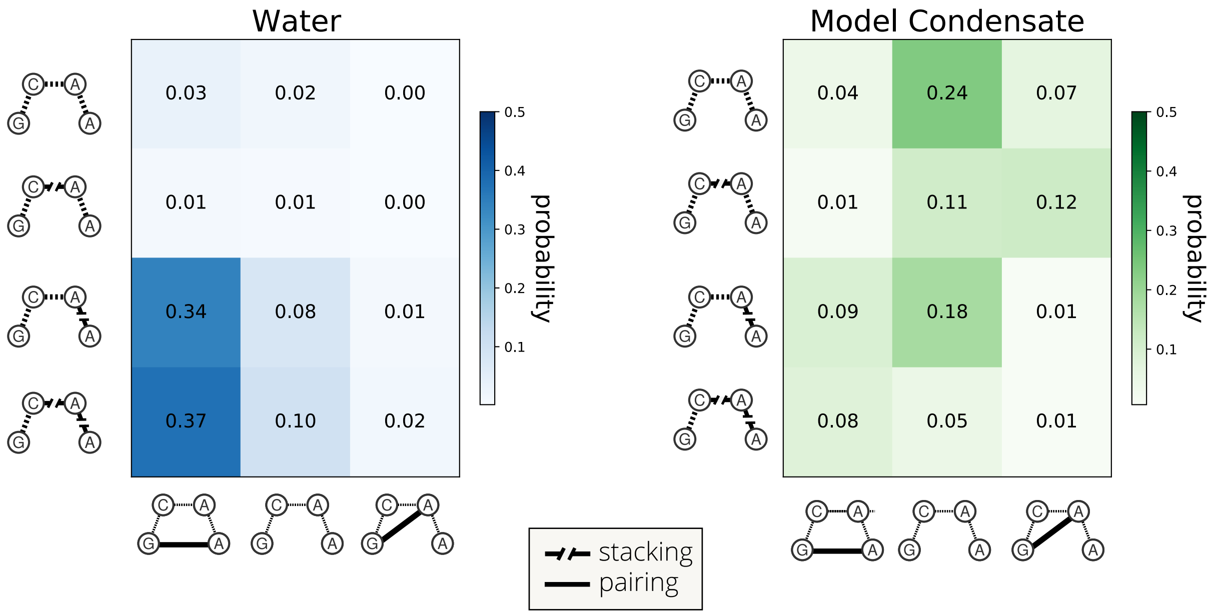


**Figure S6.** GCAA loop conformational states. Probability maps of main loop conformational states in terms of pairing and stacking for structures with fully formed stem ($Nc = 3$). In row, states are distinguished in terms of stacking (from top to bottom: no stacking, C5-A6 stacking, A6-A7 stacking, and C5-A6 stacking + A6-A7 stacking); in column, states are distinguished in terms of pairing (from left to right: G4-A7 pairing, no pairing, G4-A6 pairing). Left panel (blue): probabilities of RNA in water. Right panel (green): probabilities of RNA inside the condensed model.

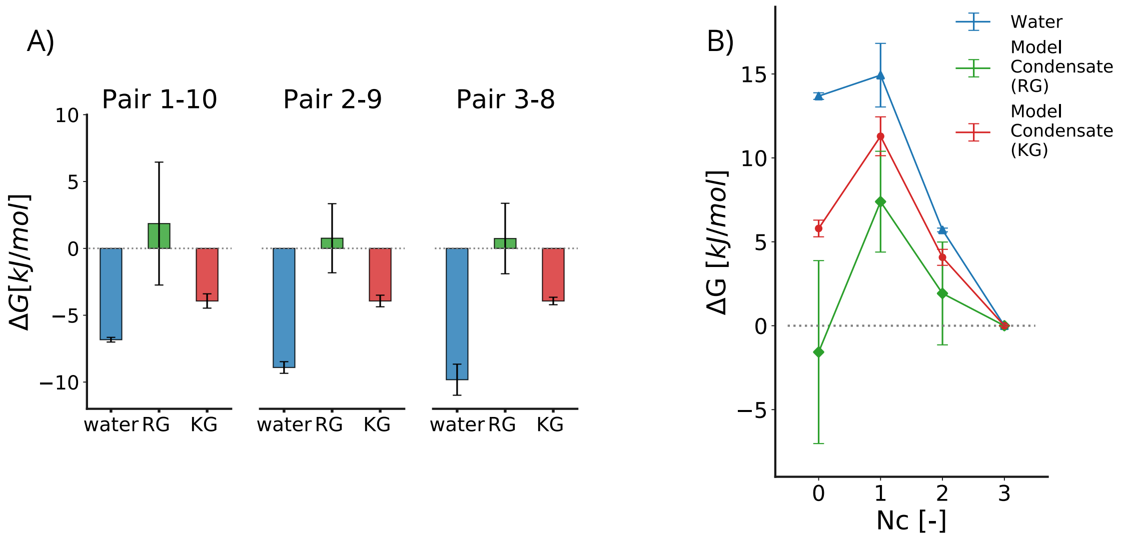


**Figure S7.** (**A**) Formation free energies ($\Delta G$) of individual pairs (G1-C10, G2-C9, C3-G8, respectively). In blue are the values obtained from the simulation of the RNA in water, in green are the values for the RNA within the model condensate consisting of a solution of RGRGG peptides (indicated RG in the figure) and in red within KGKGG peptide solution (indicated KG in the figure) . (**B**) Free-energy ($\Delta G$) as a function of the number of native stem base pairs formed ($Nc$). $Nc = 3$ indicates the fully-formed stem while $Nc = 0$ represents the absence of pairing at the stem level. The blue line indicates RNA in water, the green line refers to RNA within the argine-rich model condensate (RG) and the red line refers to RNA within the lysine-rich model condensate (KG) .

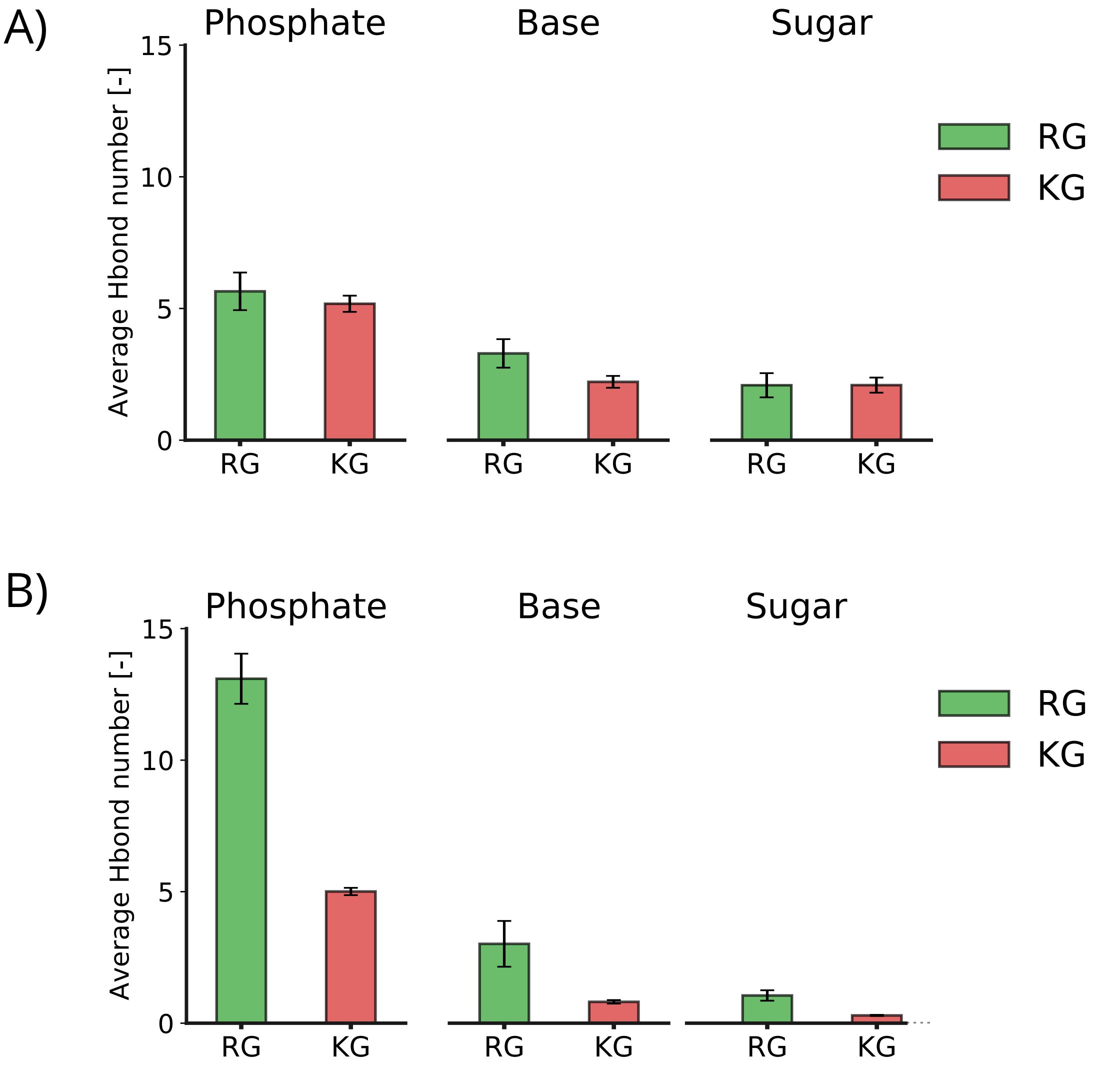


**Figure S8.** Hydrogen bond interactions. (**A**) Average hydrogen bond number between RNA and peptide main chains, reported for different RNA functional groups: phosphate (left), base (center), sugar (right). (**B**) Average hydrogen bond number between RNA and peptide side chains, reported for different RNA functional groups: phosphate (left), base (center), sugar (right).
